## Supplementary material for "Uncovering the hidden diversity and functional roles of root endophytic *Streptomyces* under drought stress": S1 Text. Supplementary figures A-G

A B

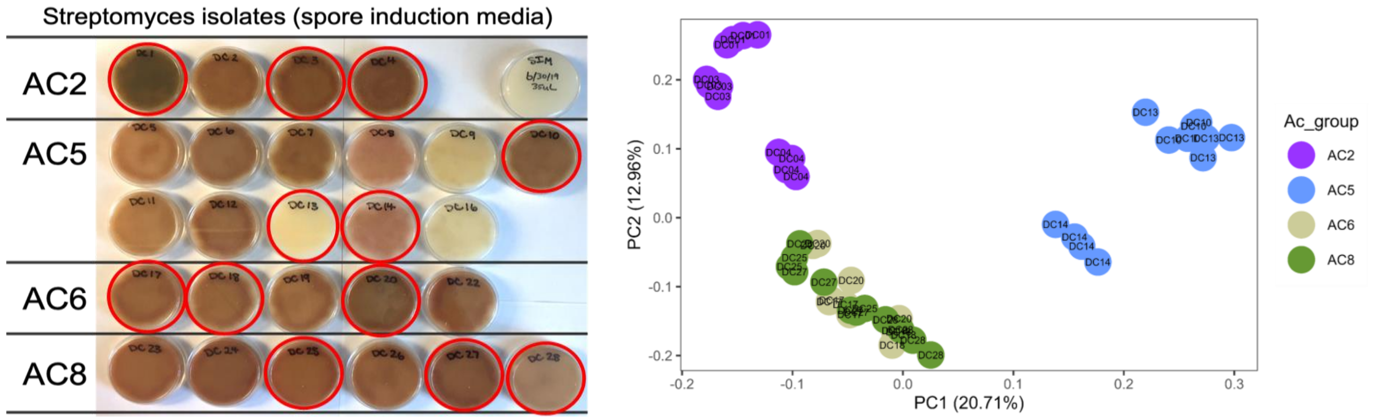


**Fig A. Phenotypic and metabolomic differentiation among *Streptomyces* isolates sharing a dominant V3-V4 16S rRNA ASV. (A)** Image of the growth (on spore induction media, SIM) for all 28 *Streptomyces* isolates with 100% identity to the dominant ASV identified in the root via V3-V4 16S rRNA sequencing with original hand labeling from Figure 1A. (**B**) Ordination of non-polar exometabolomic profiling of the 12 strains following growth on liquid tap water–yeast extract (TWYE) medium showing distinct clustering of AC group 2 and 5 (replication n=4). AC groups are presented by colors: AC2, purple; AC5, light blue; AC6, green mist; AC8, green. *Streptomyces* isolates ID are indicated in the corresponding plate.

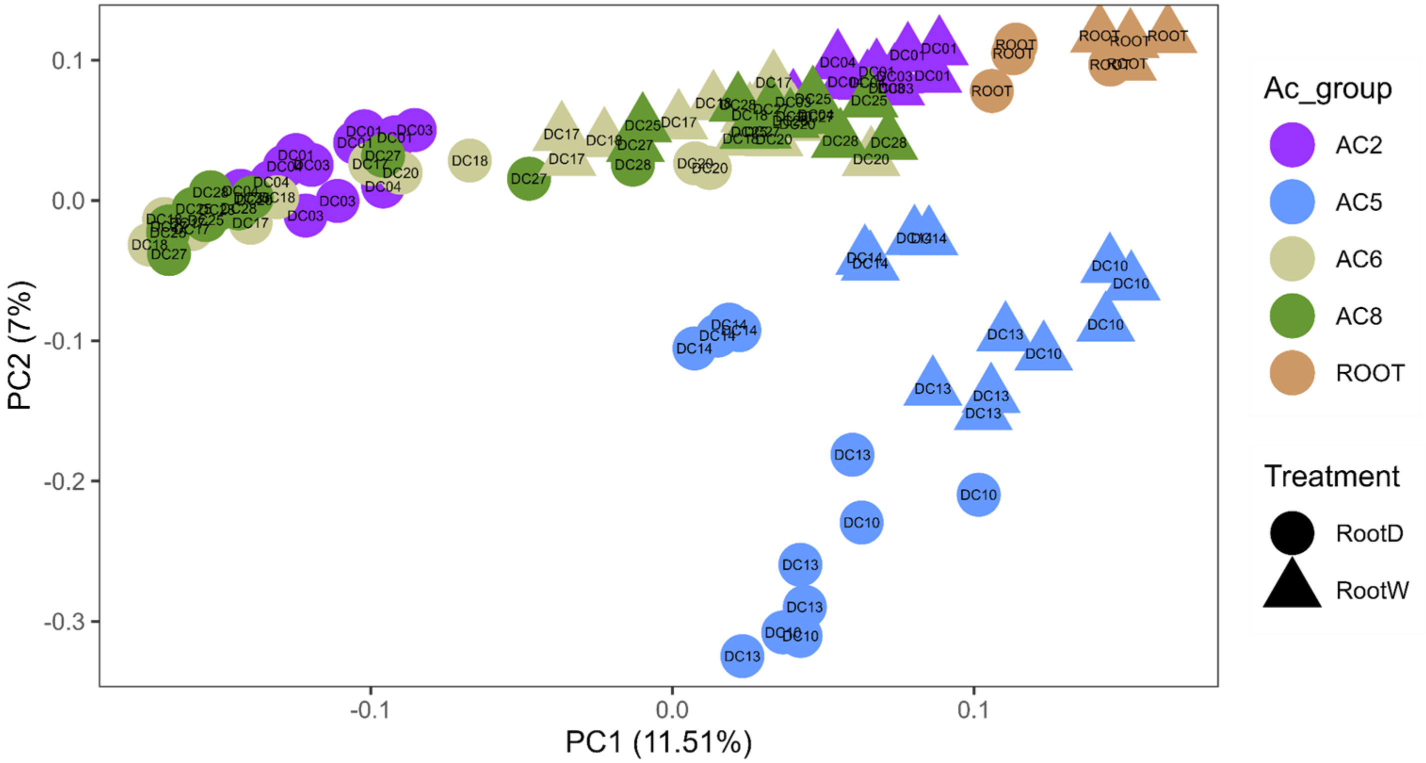


**Fig B. Ordination plot of nonpolar exometabolomics data analyzed from the spent media of all identified *Streptomyces* strains matching V3-V4 ASV in the isolate collection following growth on root tissue.** The color of each shape indicates the AC group it belongs to: AC2, purple; AC5, light blue; AC6, green mist; AC8, green. Blank control samples containing only drought root tissue (brown circles) or control irrigated root tissue (brown triangles) are shown at top right in the plot (replication n=4). *Streptomyces* isolates ID are indicated in the corresponding plate.

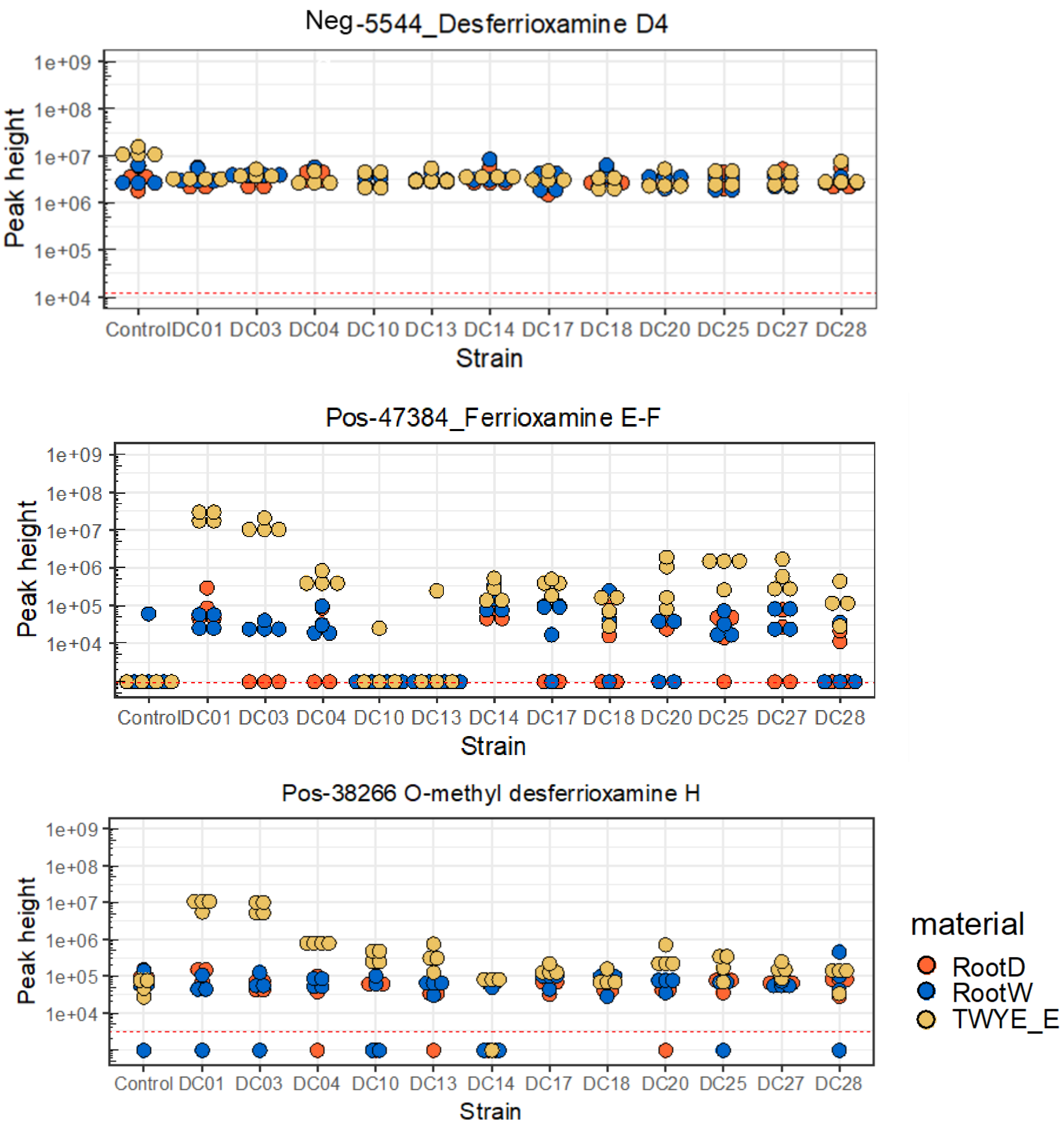


**Fig C. Production of representative individual siderophores putatively identified and measured by exometabolomics.** DC strains were under growth on TWYE (yellow), drought-stressed root tissue (orange), or non-stressed root tissue (blue). Cells were pelleted by centrifugation and the clarified supernatants were then snap-frozen in liquid nitrogen and lyophilized to dryness. Lyophilized supernatants were resuspended with methanol, sonicated and then transferred to LC-MS glass autosampler vials for untargeted liquid chromatography–mass spectrometry identification.

### MS/MS spectra mirror match

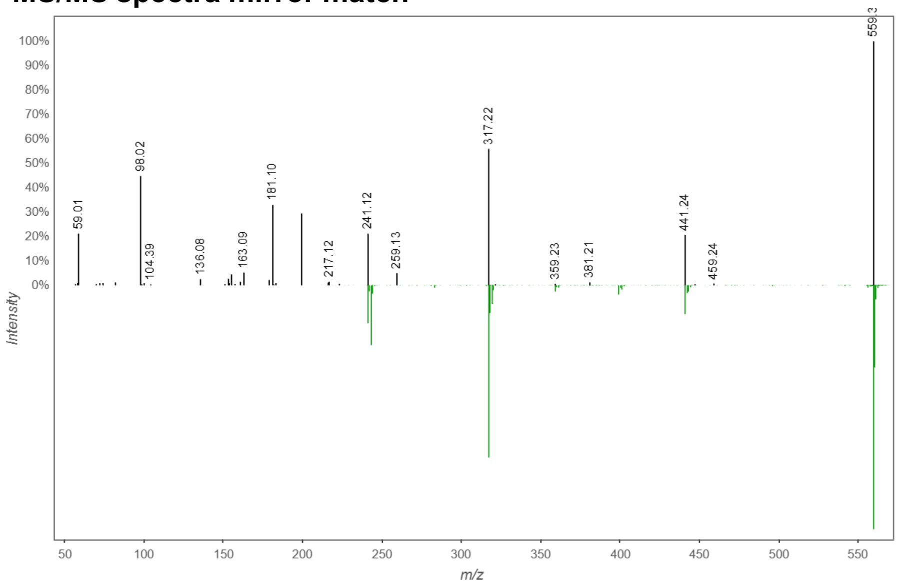


| Library ID | desferrioxamine D4 |
|---|---|
| MQScore | 0.474601 |
| Shared Peaks | 4 |
| SpecMZ | 559.35 |
| LibMZ | 559.35 |
| RTConsensus | 11.4541 |
| IonMode | Negative |
| cluster index | 5544 |
| Smiles | O=C(C)N(O)CCCCCNC(CCC(N(O)CCCCCNC(CCC(NCCCCNC([H])=O)=O)=O)=O)=O |
| SpectrumID | CCMSLIB00005724367 |
| Structure | 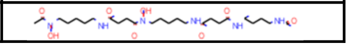 |

### MS/MS spectra mirror match

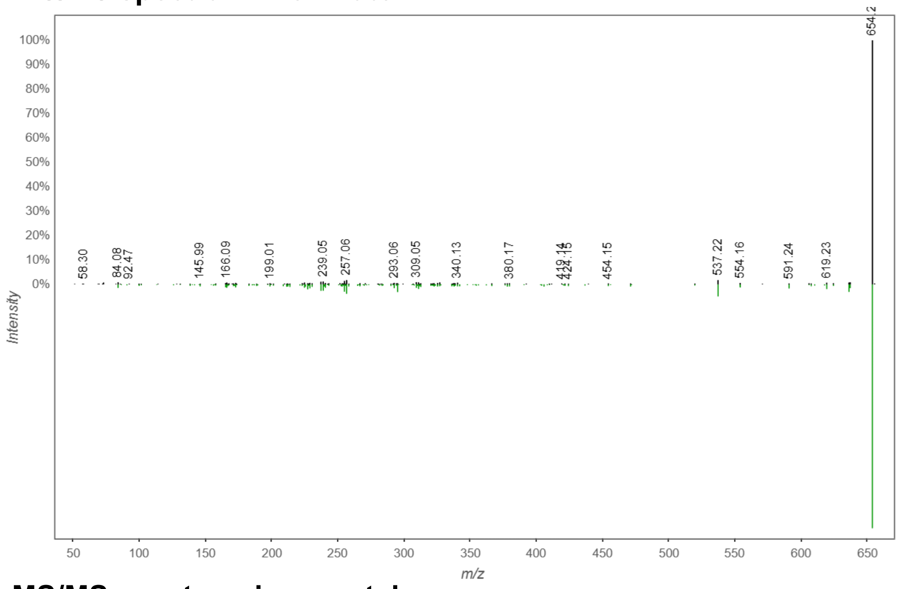


| Library ID | Ferrioxamine-E-Fe-adduct |
|---|---|
| MQScore | 0.747154 |
| Shared Peaks | 26 |
| SpecMZ | 654.26 |
| LibMZ | 654.27 |
| RTConsensus | 3.1318 |
| IonMode | Positive |
| cluster index | 47384 |
| Smiles | |
| SpectrumID | CCMSLIB00005723618 |

### MS/MS spectra mirror match

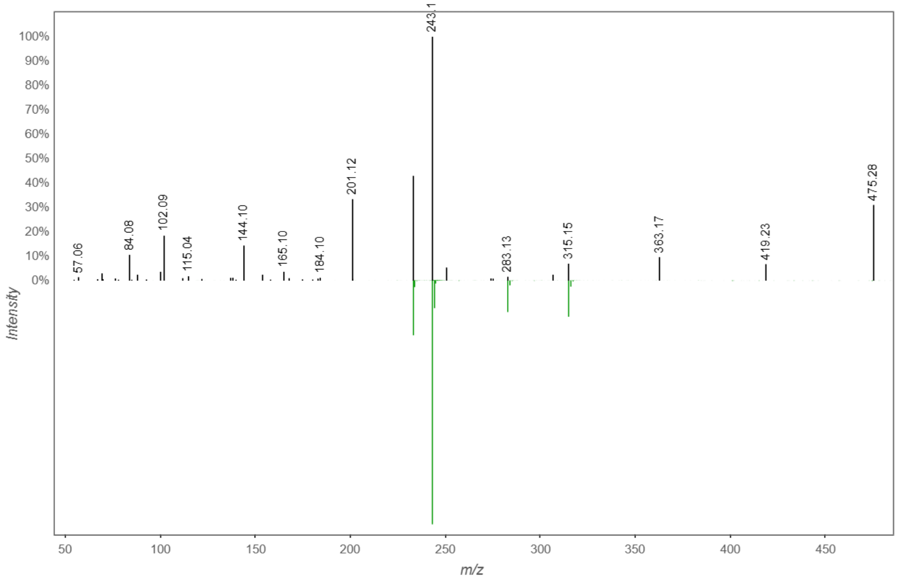


| Library ID | O-methyl desferrioxamine H |
|---|---|
| MQScore | 0.684794 |
| Shared Peaks | 4 |
| SpecMZ | 475.27 |
| LibMZ | 475.27 |
| RTConsensus | 0.9271 |
| IonMode | Positive |
| cluster index | 38266 |
| Smiles | COC(CCC(NCCCCCN(C(CCC(NCCCCN(C(C)=O)O)=O)=O)C)=O)=O.[OH] |
| SpectrumID | CCMSLIB00005723631 |
| Structure | 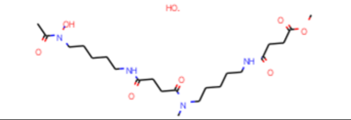 |

**Fig D. Mirror match plot comparing experimental (query) and library MS/MS spectra of siderophores putatively identified by exometabolomics.** The mirror plot visualizes the spectral alignment between an experimental MS/MS spectrum (top in black) and a reference spectrum from the GNPS library (bottom in green). Shared fragment ions are indicated by aligned peaks, and spectral similarity is quantified using the cosine score. The precursor ion m/z, retention time (RT), ionization mode, cosine similarity, and number of matched fragment peaks are used to assess the quality and confidence of the match.

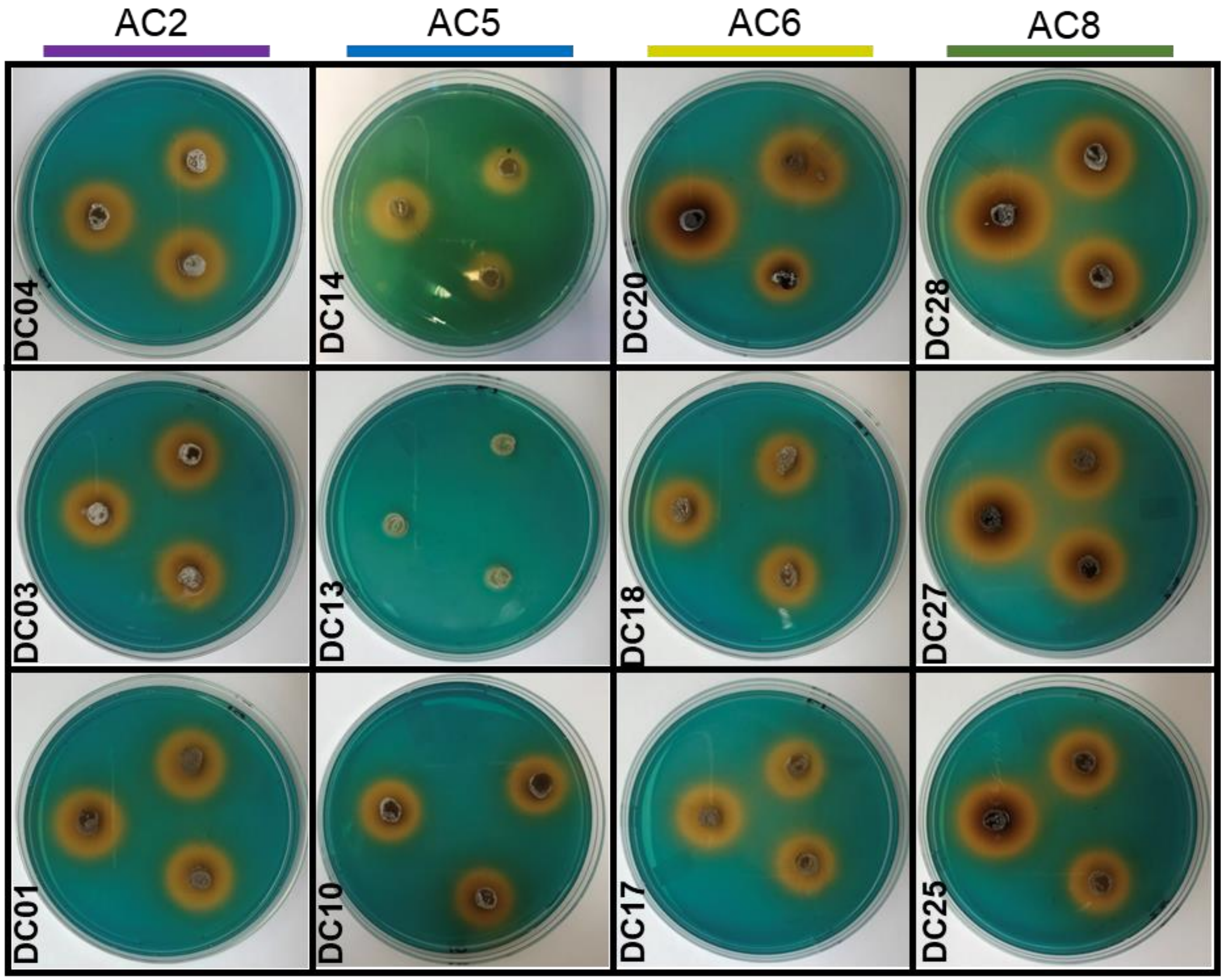


**Fig E. Total siderophore production as measured by CAS-LB agar assay based on the method of Schwyn and Neilands (1987).** *Streptomyces* isolates were spot-inoculated onto the CAS-LB plates and incubated at 28°C for 72 hours. A yellow to orange halo around colonies indicated siderophore production, resulting from iron chelation from the blue CAS–$Fe^{3+}$ complex. The experiment was repeated twice with three technical replicates each.

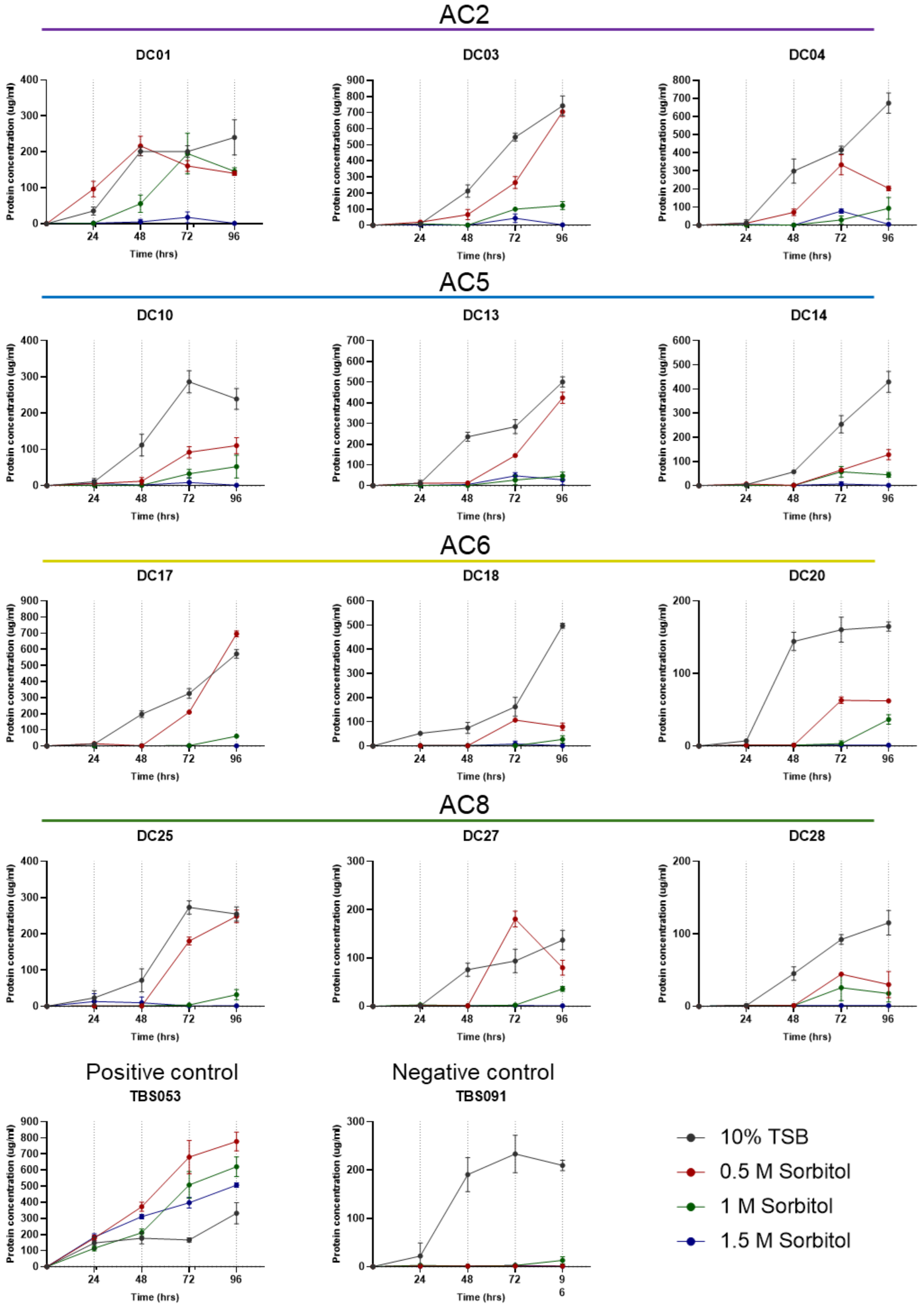


**Fig F. Osmotic tolerance assay of Streptomyces isolates under increasing sorbitol-induced stress.** *Streptomyces* isolates were grown in tryptic soy broth (10% TSB) supplemented with sorbitol at final concentrations of 0.5 M, 1.0 M, and 1.5 M. Bacterial growth was monitored over time (0, 24, 48, 72, and 96 hours; n = 3) using Bradford protein assays to quantify total biomass. This time-course approach enabled the assessment of growth dynamics and adaptation to osmotic stress. *Bacillus megaterium* (TBS 053) and *Paenibacillus lautus* (TBS 091) were included as positive and negative controls, respectively.

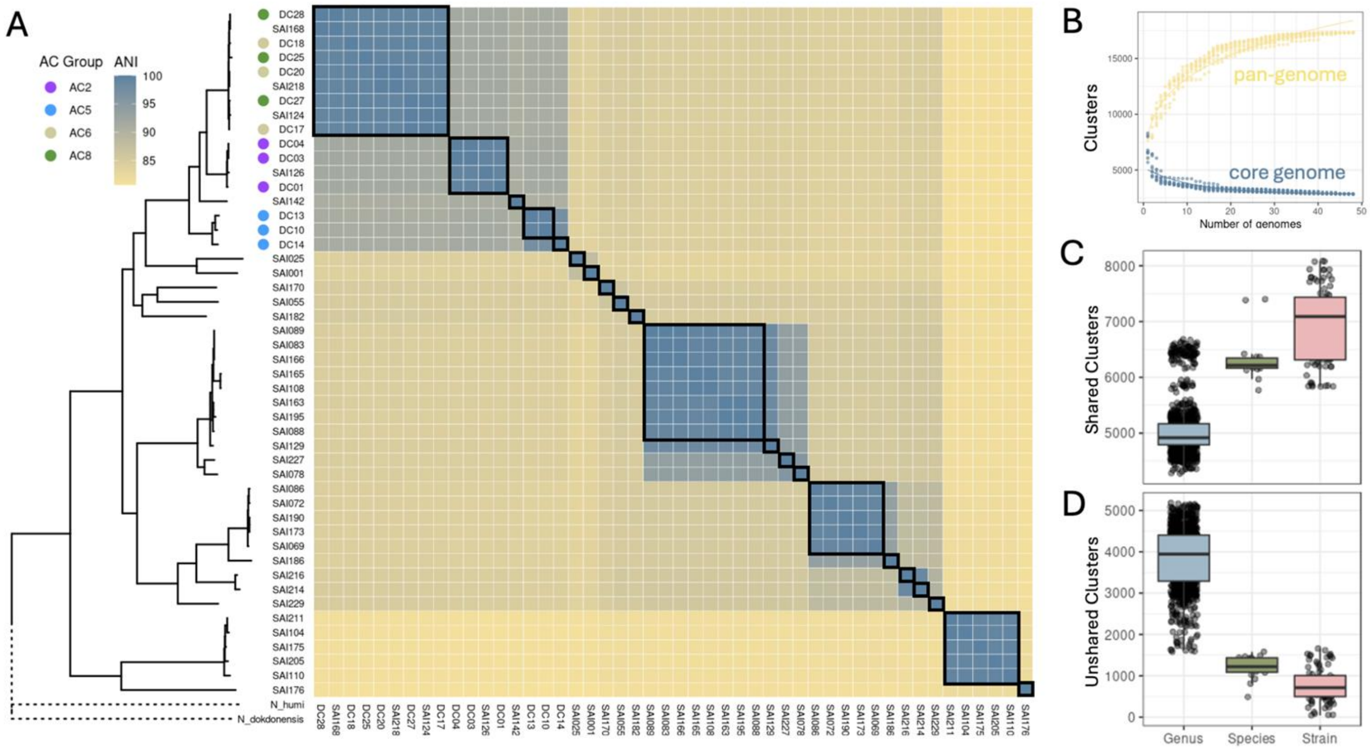


**Fig G. Phylogenomic and comparative genomic analysis of 48 *Streptomyces* isolates. (A)** Phylogenetic tree of all 48 *Streptomyces* isolates based on alignment of 138 single copy genes and rooted by a *Nocardioides* outgroup. Relevant isolates are annotated with AC Group. Corresponding ANI heatmap aligned to tree tips shows levels of relatedness among isolates calculated with *fastANI*. Black boxes within the heatmap indicate unique strains (>99% ANI). **(B)** Pan- and core genome rarefaction simulations fit to power law and exponential decay curves, respectively using *pagoo* R package. **(C)** Shared and **(D)** unshared gene cluster counts between pairs of isolates compared at different taxonomic levels across the collection. Gene clusters identified by *Orthofinder* and taxonomic classification determined by pairwise ANI (Genus: <95%, Species: 95-99%, Strain: >99%).
